# *optix* regulates abdominal melanin pigmentation in the tobacco hawkmoth *Manduca sexta*

**DOI:** 10.64898/2026.08.20.746068

**Authors:** Martik Chatterjee, Gabriel C. Hatto, Christophe Duplais, Jarrod Varnell, Robert A. Raguso, Robert D. Reed

## Abstract

Research on butterflies has uncovered a conserved “toolkit” of genes for color pattern development and evolution. One of these genes is *optix*, a homeobox transcription factor that regulates ommochrome and melanin pigmentation, as well as structural coloration, in nymphalid butterflies. It remains unclear, however, whether *optix* plays any roles in color patterning outside of the Nymphalidae. We used CRISPR-Cas9 to disrupt *optix* in the tobacco hornwormmoth *Manduca sexta* and observed a dramatic abdominal pigmentation phenotype, where orange pigmentation was replaced by black eumelanin. Chemical assays suggest that the orange pigment is not an ommochrome, indicating that *optix* modulates an alternative, uncharacterized pigment pathway in *M. sexta*. RNA-seq and chemical analyses of orange and black abdominal scales lead us to speculate that the orange pigment may be a type of melanin, perhaps N-β-alanyldopamine (NBAD) sclerotin. Our results suggest that *optix* plays a deeply ancestral role in pigment regulation in Lepidoptera, and demonstrates evolutionary flexibility in how it interfaces with pigment chemistry across moths and butterflies.

**Graphical Abstract:** 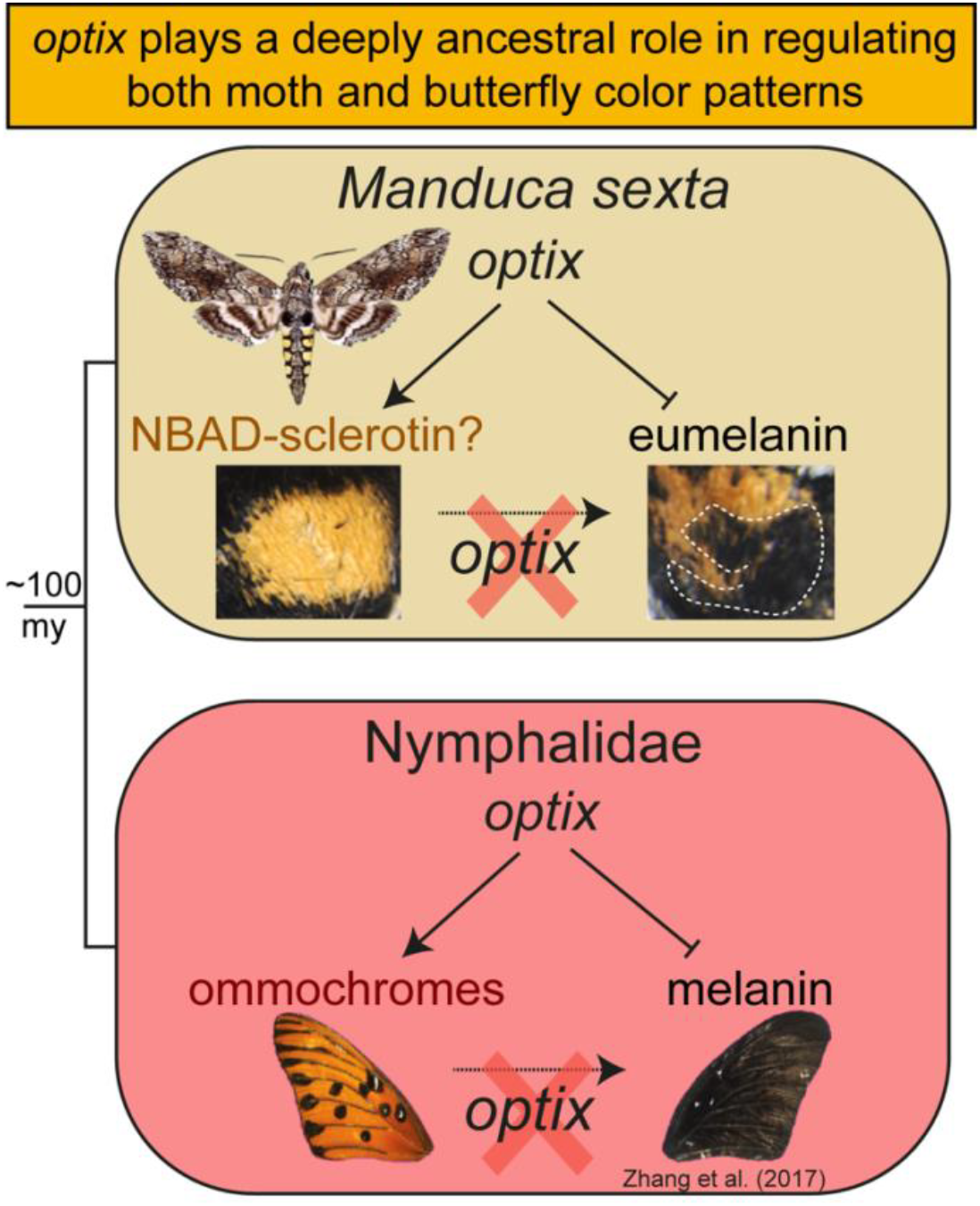

## Introduction

Genetic studies on coloration in Lepidoptera have focused largely on butterfly wing patterns, leaving other aspects of lepidopteran pigmentation comparatively understudied. Studies on moth coloration are especially lacking, despite a few notable case studies such as adaptive melanism in the British peppered moth *Biston betularia* (Hof et al., 2016), color polymorphism in the wood tiger moth *Arctia plantaginis* (Brien et al., 2023), and natural melanic polymorphism in *Anticarsia gemmatalis* (Livraghi et al., 2026). We still have a limited understanding of how the genetic regulation of pigmentation has evolved across Lepidoptera, particularly before the split of butterflies from moths ∼100 million years ago (Kawahara et al., 2023). Studies of butterfly wing coloration have highlighted a small set of conserved “toolkit” genes that drive color diversity. Of particular interest is *optix*, a homeobox transcription factor gene, that regulates the switch between ommochrome and melanin pigments and modulates structural coloration in nymphalid butterflies (Banerjee et al., 2024; Livraghi et al., 2025; Orteu et al., 2024; Prakash et al., 2022; Zhang, Mazo-Vargas, et al., 2017). Despite its involvement in wing coloration and scale structure determination across butterflies, the role of *optix* in non-nymphalid Lepidoptera remains unexplored.

In this study we assess the color patterning role of *optix* in the tobacco hornworm moth *Manduca sexta*, which is ∼100my diverged from nymphalid butterflies (Kumar et al., 2017). *M. sexta* exhibits distinct colors, including black and brown pigments in the wings and bright orange abdominal pigmentation. We used CRISPR-Cas9 to generate *optix* mosaic knockouts and found that loss of *optix* leads to a clear shift from orange to black pigmentation in the abdomen. These results suggest that *optix* is a deeply conserved regulator of pigmentation in Lepidoptera and may play a much more ancestral role in coloration than previously suspected.

## Materials and Methods

### CRISPR-Cas9 mutagenesis of *M. sexta optix*

We identified LOC115442057 as the *optix* ortholog in the *Manduca sexta* genome assembly (JHU_Msex_v1.0; NCBI RefSeq: GCF_014839805.1) (Gershman et al., 2021). We designed two sgRNAs targeting the exon containing the conserved SIX1 and homeobox domains (Figure 1b): 5′-CTCCTGGGACGAGTCCACGA-3′ and 5′-CCCCACGAAGAAACGTGAGT-3′. We screened for off-targets by BLAST against the *M. sexta* genome and detected no significant matches outside the *optix* locus. We rehydrated sgRNAs (IDT) in 1× Tris–EDTA and complexed them with Cas9 (IDT) to final concentrations of 150–250 ng/µL sgRNA and 300–500 ng/µL Cas9. We collected eggs laid by mated female moths on leaves of *Datura wrightii* (1–2 h into scotophase), rinsed and mounted them on glass slides, injected embryos within 4 h with the sgRNA–Cas9 mix, then reared larvae on artificial diet (27°C, 65% RH, 16 h:8 h L:D) before freezing adults (™20°C) for imaging.

**Figure 1.**
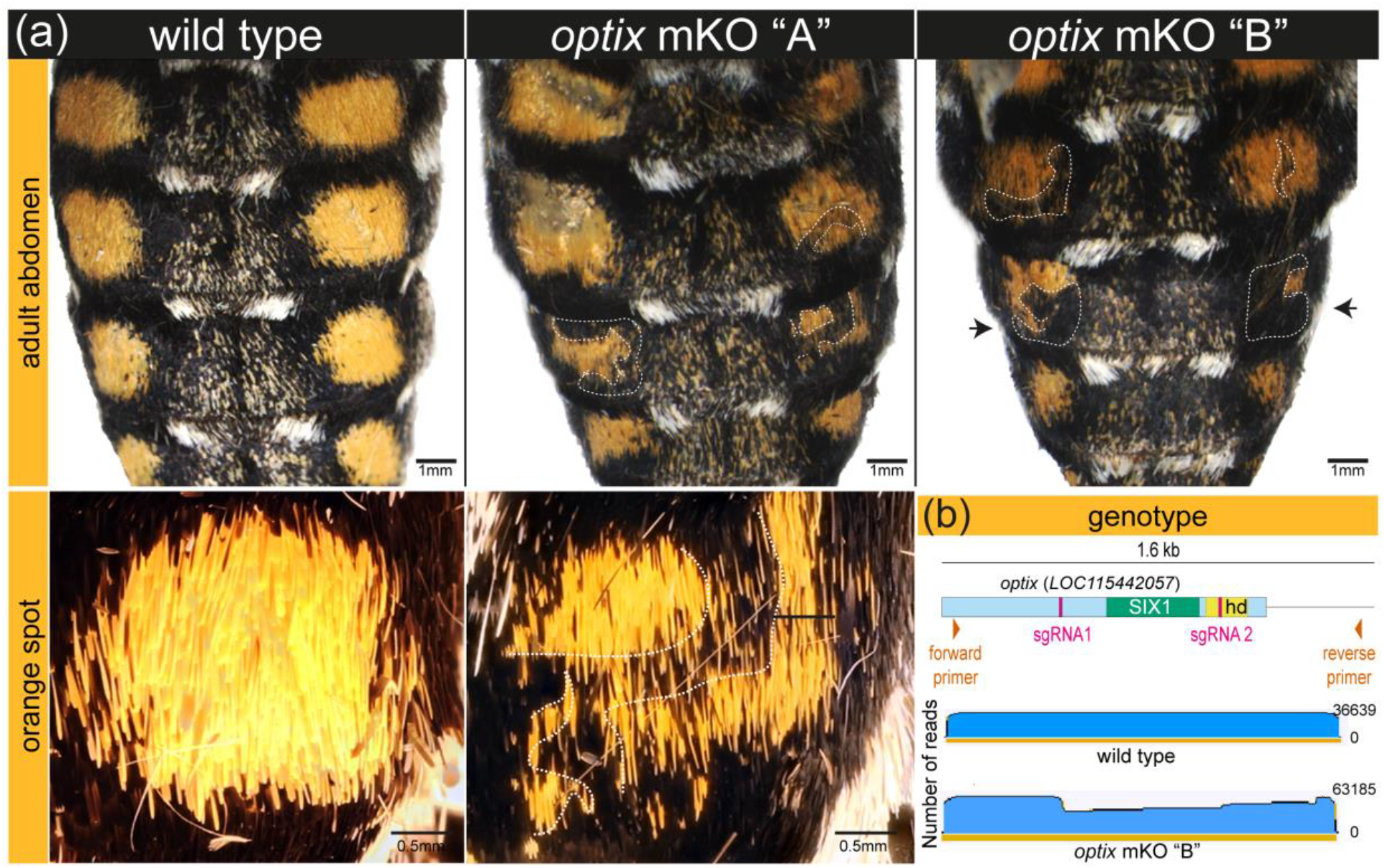
CRISPR-Cas9-mediated mosaic knockout (mKO) of *optix* switches abdominal pigmentation from orange to black in *M. sexta*. (a) Wild-type individual (*left*) compared with two *optix* mKO mutants, “A” (*middle*) and “B” (*right*), showing black mosaic patches (*outlined with dashed white lines*) in the otherwise orange abdominal segments. Bottom panels show magnified views of corresponding segments from the wild type and mKO “B.” A black arrow highlights the near-complete, asymmetrical transformation of an orange region to black in mKO “B.”(b) Genotyping of mKO “B” confirms a deletion at the *optix* locus at the sgRNA1 cut site, visualized by read pileups (blue). hd: homeobox domain. For detailed sequence alignments, see S.I. Figure 1.

To distinguish mortality caused by embryo manipulation from mortality associated with the *optix-*targeting sgRNA, we performed parallel water- and Cas9-only injection controls and scored hatching rate. We also screened unmanipulated wild-type adults (n=40) drawn from multiple laboratory colonies for spontaneous abdominal color variants resembling the CRISPR phenotypes.

### Genotyping *M. sexta optix* phenotypes

To validate CRISPR-induced lesions, we extracted genomic DNA from the abdominal segment of an individual with a clonal black patch and from a wild-type control (OMEGA BIO-TEK). We PCR-amplified a region spanning the predicted cut sites using primers 5′-GATAGAGGTAGGCGGCGGT-3′ and 5′-AACCGATACACTGGACACGC-3′, purified amplicons with AMPure XP beads (0.9×), and sequenced them on an Oxford Nanopore MiniION. We aligned reads to the *optix* scaffold with minimap2 (-x map-ont) (Li, 2018) and inspected indels at the target site in Geneious Prime 2023.1.2.

### Characterization of pigment types

As an initial assay for ommochrome pigments, we extracted scales from orange abdominal segments of *M. sexta* and from red wing patterns of *Heliconius sp*. (positive controls) in 1,000 µL of 1% HCl in methanol (v/v) for 1hr at room temperature, filtered extracts, and measured UV–Vis absorbance on a Genesys 10S spectrophotometer from 190–800 nm at 5-nm intervals.

To more broadly assay for ommochrome, pterin, and melanin pigments we analyzed extracts of orange and black abdominal scale patches by LC–HRMS, melanin degradation–LC–HRMS, and Raman spectroscopy. For LC–HRMS, we suspended orange dorsal abdominal segments of *M. sexta* in 0.5 mL methanol containing 0.1% formic acid, homogenized tissue with a Bead Ruptor (Omni International, Kennesaw, GA, USA) using stainless steel beads, filtered homogenates through 0.2-µm PTFE filter vials, and transferred filtrates to 200-µL glass insert vials for analysis. We used pterin (xanthopterin) and ommochrome (3-hydroxykynurenine and xanthurenic acid) standards to detect and quantify putative pigments. Because pure standards were unavailable for xanthommatin, hydroxyxanthommatin, and rhodommatin (ommochromes) and for chrysopterin and erythropterin (pterins), we used *Danaus plexippus* wing and *Oncopeltus fasciatus* exoskeleton extracts to identify and quantify these compounds.

After ruling out ommochromes and pterins, we tested for melanins (pheomelanin and eumelanin) using a modified protocol of Ito et al. 2011 (Affenzeller et al., 2019; Ito & Wakamatsu, 2011) where melanin degradation products are quantified through an oxidative reaction with H_2_O_2_. We suspended 6 mg of black or orange dorsal abdominal scales in 100 µL deionized water (2-mL tube), added 375 µL of 1 M K_2_CO_3_ and 40 µL of 30% H_2_O_2_, and rotated samples at 25°C for 24 h. We quenched residual H_2_O_2_ with 50 µL of 10% Na_2_SO_3_, acidified with 150 µL of 6 M HCl, centrifuged at 5,000 g for 1 min, and collected a 10-µL aliquot. We analyzed extracts on an Agilent 1260 LC with a C18 column (21.2 × 150 mm, 5 µm) using a methanol–water gradient containing 0.1% formic acid, and quantified eumelanin and phenomelanin degradation products– pyrrole-2,3-dicarboxylic acid (PDCA) and thiazole-4,5-dicarboxylic acid (TDCA), respectively—by comparing peak areas to injections of pure standards purchased from Sigma-Aldrich.

We acquired data on an Agilent 6545 qTOF HRMS in negative ion mode, processed results in Agilent MassHunter Quantitative Analysis (v10.1), and reported PDCA and TDCA concentrations in µg·g^-1^ tissue.

We collected Raman spectra of the abdominal scale pigments on a WITec Alpha300R confocal Raman microscope (100x, 0.9 NA objective; 785-nm excitation at 0.4 mW; 300 g/mm diffraction grating), recording for 40 s at three locations per sample and averaging them. We applied baseline subtraction, normalization, and curve smoothing (to remove instrument-related interference) and compared spectroscopic bands to published eumelanin and pheomelanin values (Galván et al., 2013).

### Transcriptomic profile of abdominal segments

We dissected pupae at ∼75% of pupal development, when black abdominal pigmentation is established, and collected tissue from the black dorsal midline, black abdominal segments, and future orange abdominal segments (minimal/no visible pigment at this stage. We stored tissue in TRIzol™ at ™80°C, extracted RNA with the TRIzol™ Plus kit, and sequenced libraries with RQN > 7 (Novogene; 150-bp paired-end). For each replicate, we pooled tissue from 2–3 individuals and generated four replicates for black abdomen and dorsal midline and three replicates for future orange abdomen.

We aligned reads to the *M. sexta* genome with STAR and quantified gene-level counts using --quantMode GeneCounts. We performed differential expression analysis in DESeq2 (Wald test) and defined DEGs as padj < 0.05 and |log2FC| > 1. We report gene names for annotated orthologs in the main text and provide gene model IDs and names for all DEGs in Supplementary File 1.

## Results and Discussion

### optix is as a regulatory switch between orange and black pigments in the M. sexta

#### abdomen

Because CRISPR-Cas9 injections in early syncytial embryos typically produce mosaic knockouts (mKOs) affecting only a subset of cell lineages, and because disrupting a pleiotropic homeobox gene such as *optix* can reduce viability, we expected color phenotypes in only a small proportion of surviving adults. Of 670 eggs injected with *optix* sgRNA-Cas9 complexes, 92 hatched (13.7%) and 33 reached adulthood (4.9% of injected eggs; 35.9% of hatchlings); two of the 33 adults (6.1%) showed clear mosaic color phenotypes (Table 1). Both mutants were recovered from the lowest sgRNA concentration tested (150 ng/µL), which also produced 29 of the 33 surviving adults. By contrast, approximately 70% of embryos hatched in both the water-only and Cas9-only control treatments (Table S1), indicating that the low hatch rate was not explained by microinjection or Cas9 alone. Low recovery is also consistent with published nymphalid *optix* CRISPR screens, which likewise reported low hatch and adult-recovery rates (Zhang, Mazo-Vargas, et al., 2017).

**Table 1.** *optix* CRISPR-cas9 microinjection outcome.

| **sg RNA conc.**<br>**(ng/μl)** | **Eggs injected** | **Hatched** | **Eclosed** | **Mutants** |
| --- | --- | --- | --- | --- |
| 250 | 150 | 4 | 0 | 0 |
| 200 | 121 | 21 | 4 | 0 |
| 150 | 399 | 67 | 29 | 2 |
| **Total** | **670** | **92** | **33** | **2** |

In both *optix* mKOs, normally orange abdominal scales contained sharply bounded clonal patches of black pigmentation (Figure 1), asymmetrically distributed across segments as expected for CRISPR-induced mosaicism (Zhang & Reed, 2017). In one individual (mKO “B”), an entire abdominal segment shifted from orange to black.

Genotyping of black tissue confirmed indels at the sgRNA target site (Figure 1b; Supplementary Figure 1). Unlike *optix* knockouts in nymphalid butterflies, surviving *M. sexta* adults showed no obvious wing phenotypes, although limited survival and body-restricted mosaicism could have masked wing effects. Importantly, we did not observe comparable clonal patches of black scales within orange patterned regions in the surviving control- and the randomly sampled-adults (chi-squared test, *p < 0*.*005*), supporting the conclusion that the mosaic pigmentation phenotypes recovered in the two mutants resulted from CRISPR-induced disruption of *optix*.

The abdominal orange-to-black switch we observed in *M. sexta* resembled *optix* knockout phenotypes in nymphalid wings where *optix* promotes orange and red ommochromes, while repressing melanins (Zhang, Mazo-Vargas, et al., 2017). We therefore sought to test if orange abdominal pigments in *M. sexta* are ommochromes, which would be consistent with the function of *optix* in butterflies. We found that orange abdominal pigments of *M. sexta* were insoluble in acidified methanol, which readily extracts ommochromes from insect tissues, including butterfly scales (Linzen, 1974), and we detected no ommochrome-like UV–Vis absorbance profiles in extracts (Figure S2). Additionally, chemical analysis using LC-HRMS similarly failed to detect ommochromes (xanthommatin, hydroxyxanthommatin, rhodommatin) or pterins (xanthopterin, chrysopterin, erythropterin). We detected only trace 3-hydroxykynurenine (<10 µg/g), far below levels typically associated with visible pigmentation which generally occur at mg/g levels. Thus, it appears that ommochromes or pterins do not contribute to orange abdominal pigmentation in *M. sexta*.

Because the orange abdominal pigment proved insoluble in standard solvents, we hypothesized that it may be a type of melanin, potentially pheomelanin given its orange hue and prior reports of pheomelanin-based yellow-orange coloration in some insects, including tiger moths (Brien et al., 2023; Galván et al., 2015; García et al., 2016; Hines et al., 2017; Polidori et al., 2017). Using LC-MS analysis of melanin degradation products, we detected eumelanin in black abdominal scales of *M. sexta*, but pheomelanin was undetectable from the orange abdominal scales. LC–MS of oxidative degradation products and Raman spectroscopy both corroborated these results: black abdominal scales matched characteristic eumelanin bands (Figure 2), however the distinct Raman spectrum presented by the orange scales did not match any published reference data for melanin including for pheomelanin. Given its insolubility, its yellow-brown coloration, and its distinct spectral profile, we speculate that it may instead be N-β-alanyldopamine (NBAD) sclerotin, a catecholamine-derived pigment previously implicated in cuticle tanning and yellow-to-brown pigmentation in the pupal cuticle of *M. sexta* (Hopkins et al., 1982). This interpretation remains provisional, however, because we lacked a reference Raman spectrum or other positive control for NBAD sclerotin and therefore could not independently verify the pigment’s identity.

**Figure 2.**
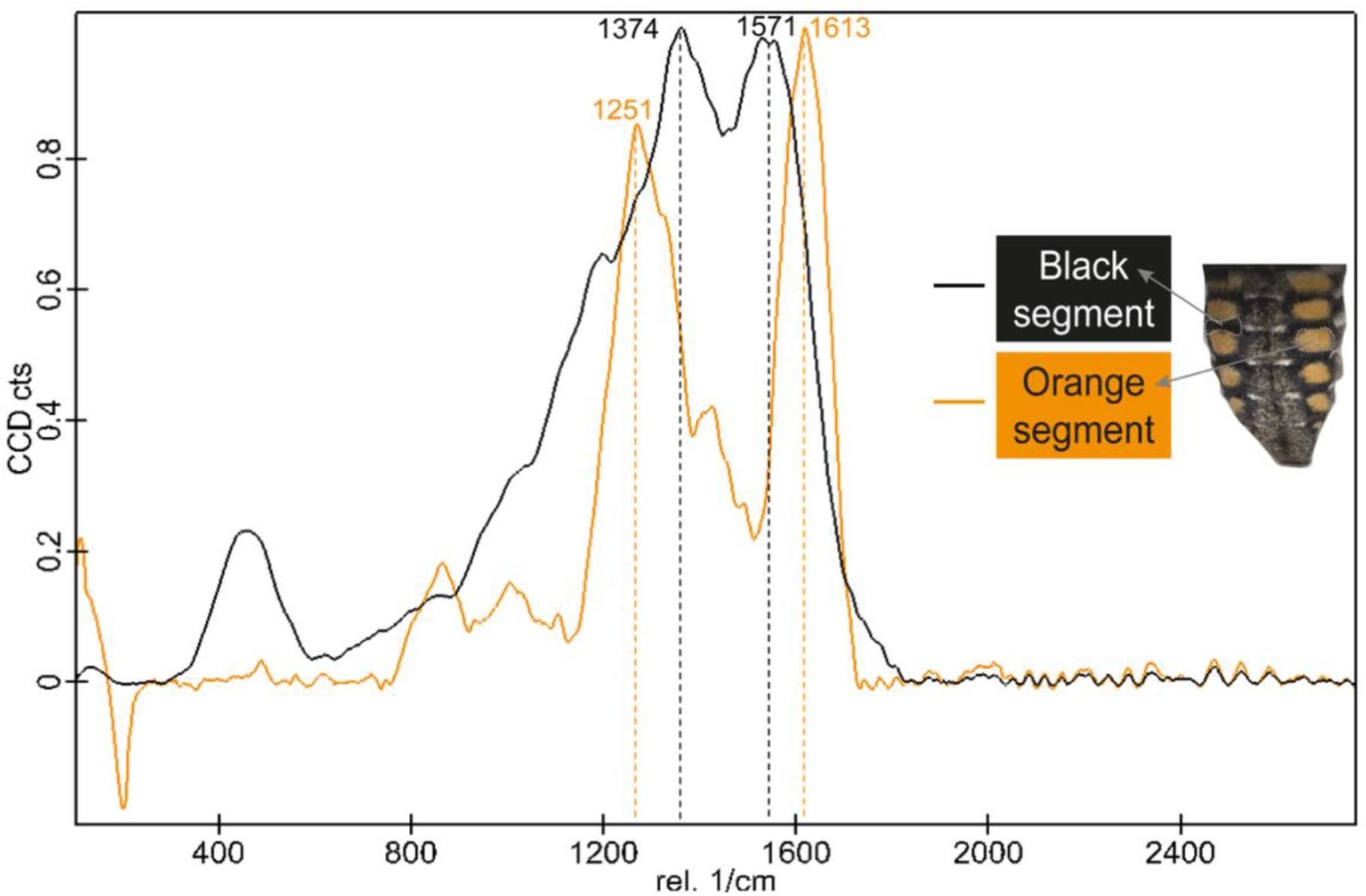
Raman spectra of abdominal pigments in *M. sexta*. Spectra were obtained by confocal Raman microscopy from the black (black line) and orange (orange line) abdominal samples. The black pigment exhibits the characteristic eumelanin signature (Galván et al., 2013), including the two peaks at ∼1380 and ∼1580 cm^-1^. In contrast, the orange abdominal pigment shows a distinct spectral profile that does not correspond to published references we are aware of, including pheomelanin (∼500, ∼1490, and ∼2000 cm^-1^).

Together, our results suggest that *optix* acts as a pigment-type switch in the abdomninal scales of *M. sexta*, promoting orange pigmentation of an unconfirmed nature, while repressing black eumelanin. To our knowledge this is some of the first evidence that *optix* plays a role in regulating pigmentation outside of nymphalid butterflies. These results suggest that a role for *optix* in pigment regulation may predate the deployment of ommochromes in butterfly wings, and prompts new questions about how the developmental genetic regulation of pigment chemistry evolved across Lepidoptera.

#### RNA-seq reveals candidate genes for abdominal pigmentation and patterning

In part because our chemistry work did not definitively identify the orange abdominal pigmentation in *M. sexta*, we performed RNA-seq to see if we could identify pigment-specific gene expression that may yield clues about its pigment type. For this work we dissected tissues from the orange and black regions of the abdominal tergum, as well as the dorsal midline, which contains a mixture of orange, black, and white scales (Figure 3). Tissues were collected at approximately 75% of pupal development—a stage when black pigments have already been deposited and orange pigments are beginning to appear. Through pairwise comparisons of gene expression between tissue types, we identified several differentially expressed genes associated with pigmentation, body segmentation, and regional patterning (Figure 3; Supplementary File 1; Wald test, padj < 0.05, |log2 fold change| > 1).

**Figure 3.**
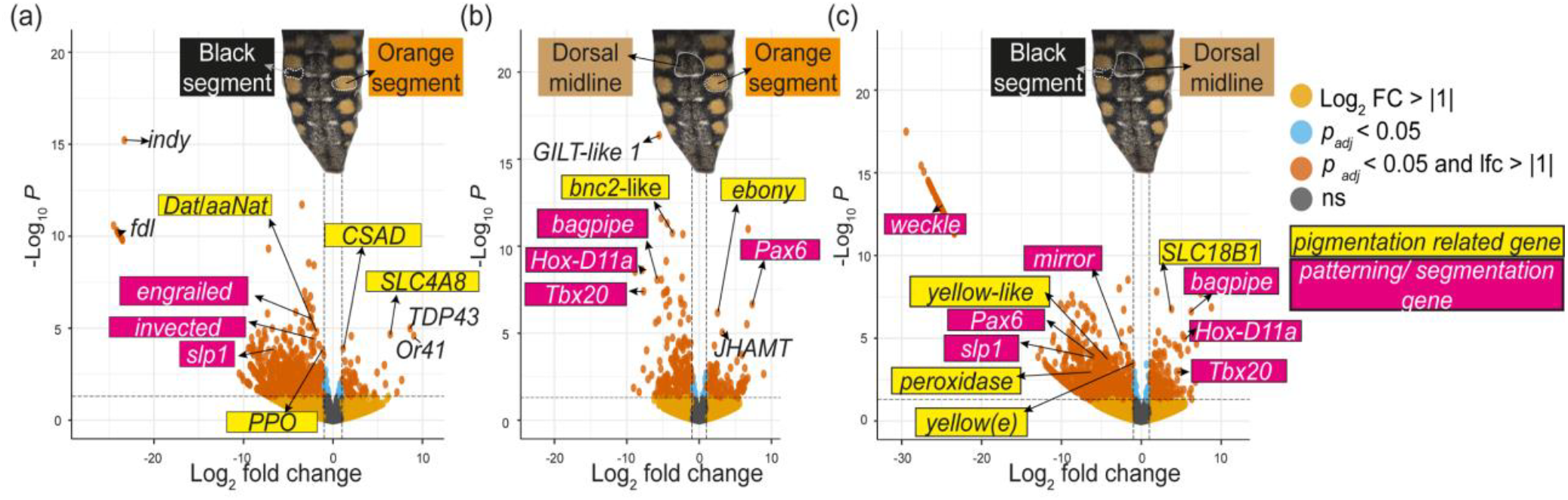
Differential gene expression across abdominal regions of *M. sexta* during late pupal development identifies candidate genes associated with pigmentation and segmental identity. Volcano plots depict pairwise comparisons of gene expression between (a) orange vs. black segments, (b) orange vs. dorsal midline, and (c) dorsal midline vs. black. Dotted lines indicate significance thresholds of adjusted *p* < 0.05 and absolute log_2_ fold change (|log_2_FC|) > 1.

In the comparison between black and orange abdominal tissues, *cysteine sulfinic acid decarboxylase* (*CSAD*, LOC115451092) was significantly upregulated in the orange region. This is notable because *CSAD* encodes an enzyme previously associated with natural melanic variation in the nymphalid butterfly *Bicyclus anynana* (Saenko et al., 2012). Similarly, *SLC4A8* (LOC119192279), a member of the major facilitator superfamily (MFS) of transporter proteins—many of which have been linked to pigmentation in insects (Zhang, Martin, et al., 2017)—also showed higher expression in the orange tissue compared to the black. In contrast, canonical melanin biosynthesis genes such as *dopamine N-acetyltransferase* (*Dat*, LOC115452451), which catalyzes the conversion of dopamine to N-acetyl dopamine (Dai et al., 2010) and *prophenoloxidase* (*PPO*, LOC115452451), a key enzyme in melanin synthesis and immune melanization (Lu et al., 2014), were significantly upregulated in the black tissues.

When we compared the orange tissue to the dorsal midline, we found that the melanin suppressor gene *ebony* (LOC115449441) was upregulated in the orange tissue. The upregulation of *ebony* in the orange tissue compared to the predominantly black dorsal midline is particularly important because Ebony activity facilitates the binding of β-alanine and dopamine to form NBAD, the precursor of yellow/orange NBAD-sclerotin throughout insects (Liu et al., 2016), adding support to our hypothesis that the orange pigment could be NBAD sclerotin. If confirmed, NBAD-sclerotin pigmentation would broaden the known range of pigment chemistries regulated by *optix* in Lepidoptera beyond the ommochromes typically associated with *optix* in nymphalid butterfly wings. This would suggest that the evolutionary conservation of *optix* as a color-pattern regulator does not necessarily depend on conservation of the downstream pigment pathway it controls. When comparing the black segments to the dorsal midline, we identified higher expression of the MFS transporter gene *SLC18B1* (LOC115445123) in the dorsal midline. Conversely, melanin-related genes such as *yellow* (LOC119193478), *yellow-e* (LOC115441510), and *peroxidase* (LOC115448945) were more strongly upregulated in the black scales. In sum, our RNA-seq works illustrates differential regulation of a number of key melanin genes across orange, black, and midline scales, including *ebony* in orange scales. Yet we did not see similar patterns for any known ommochrome or pterin genes.

Beyond pigmentation, we noted a number of patterning and segmentation genes that were differentially expressed between types and therefore may represent candidate regulators that could work in concert with *optix* to determine abdominal color patterns (Fig. 3). We did not identify any such candidates uniquely associated with the orange patch across comparisons, although the ortholog of *sloppy paired 1* (LOC115442013) was consistently associated with the black patch. We also noted that orthologs of *bagpipe* (LOC115442839), *Hox-D11a* (LOC115442839), and *Tbx20* (LOC115442839) were consistently associated with the dorsal midline across comparisons.

Together, our differential expression analysis identified a suite of candidate pigmentation genes and transcriptional regulators that may contribute to not only pigmentation but also to abdominal regional identity. These candidates, including the ones listed in Supplementary File 1, warrant functional validation to clarify their roles in pigment regulation and to test whether they act downstream of *optix* in mediating the switch between orange and black pigmentation.

## Conclusion

Our findings extend the known role of *optix* as a key color regulator beyond nymphalid butterflies into a moth lineage that diverged from the nymphalids more than 100my ago. The identity of the orange pigment in *M. sexta* adds an unexpected twist, however: unlike butterflies, where *optix* promotes ommochrome pigmentation, the chemical properties of *M. sexta* orange pigment suggest it is not an ommochrome, but possibly some other melanin derivative (e.g. NBAD sclerotin). This raises the possibility that *optix* functions as a developmental switch between catecholamine-derived pigmentation pathways in moths, potentially promoting NBAD sclerotin over eumelanin rather than toggling between ommochrome and melanin pathways as in butterfly wings. If correct, this represents a striking example of evolutionary flexibility in pigment regulatory networks – *optix* appears to maintain its high-level role in color specification while its downstream effectors have shifted dramatically across Lepidoptera. Such modularity could facilitate the diversification of color phenotypes, allowing major pigment transitions without disrupting conserved pattern formation systems. Future work aimed at biochemically confirming the orange pigment in *M. sexta* and functionally characterizing the *optix* regulatory network will be essential for understanding how pigment-type switches evolve and contribute to the spectacular diversification of insect coloration.

## Data availability

Raw sequencing reads for RNA-seq are available has been made available under BioProject ID PRJNA1444720.

## Acknowledgments

We thank Ajinkya Dahake, and Richard Fandino for their assistance in rearing *M. sexta*. We are also grateful to the Cornell University Insect Collection for their help with photographing mutant specimens. Some of this research was conducted as part of GH’s undergraduate honor’s thesis in partial fulfillment of the requirements for the Biological Sciences Honors Program at Cornell University. We are grateful to Kevin Silverstein of the Cornell Center for Materials Research shared instrumentation facility for conducting the Raman analysis. This work was supported by the United States National Science Foundation (IOS-2128164 and DEB-2242865 to RDR and DGE-2139899 to JMV), as well as a Cornell Summer Experience Grant and an Office of Undergraduate Biology Fellowship awarded to GH.

