## Supplementary material for "*optix* regulates abdominal melanin pigmentation in the tobacco hawkmoth *Manduca sexta*": S.I

**Supplementary Information**
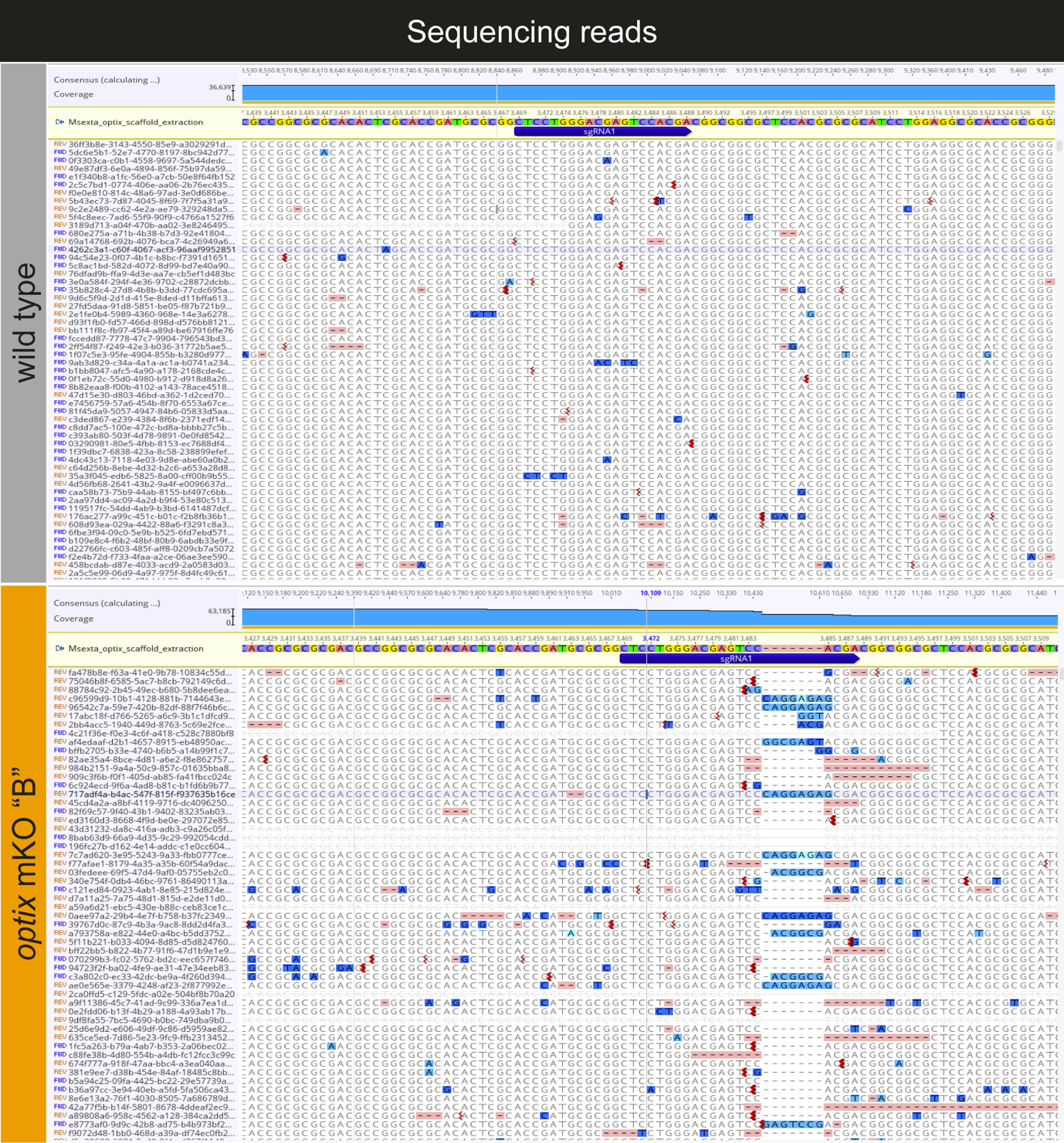

**Figure S1. *optix* sequences from mutant *M. sexta* abdominal tissues reveal deletions in the *optix* coding region.** Shown are Nanopore long reads from a wild-type individual (top) compared to the *optix* mKO “B” individual (bottom), aligned to the genome at the sgRNA1 cut site (highlighted in blue).

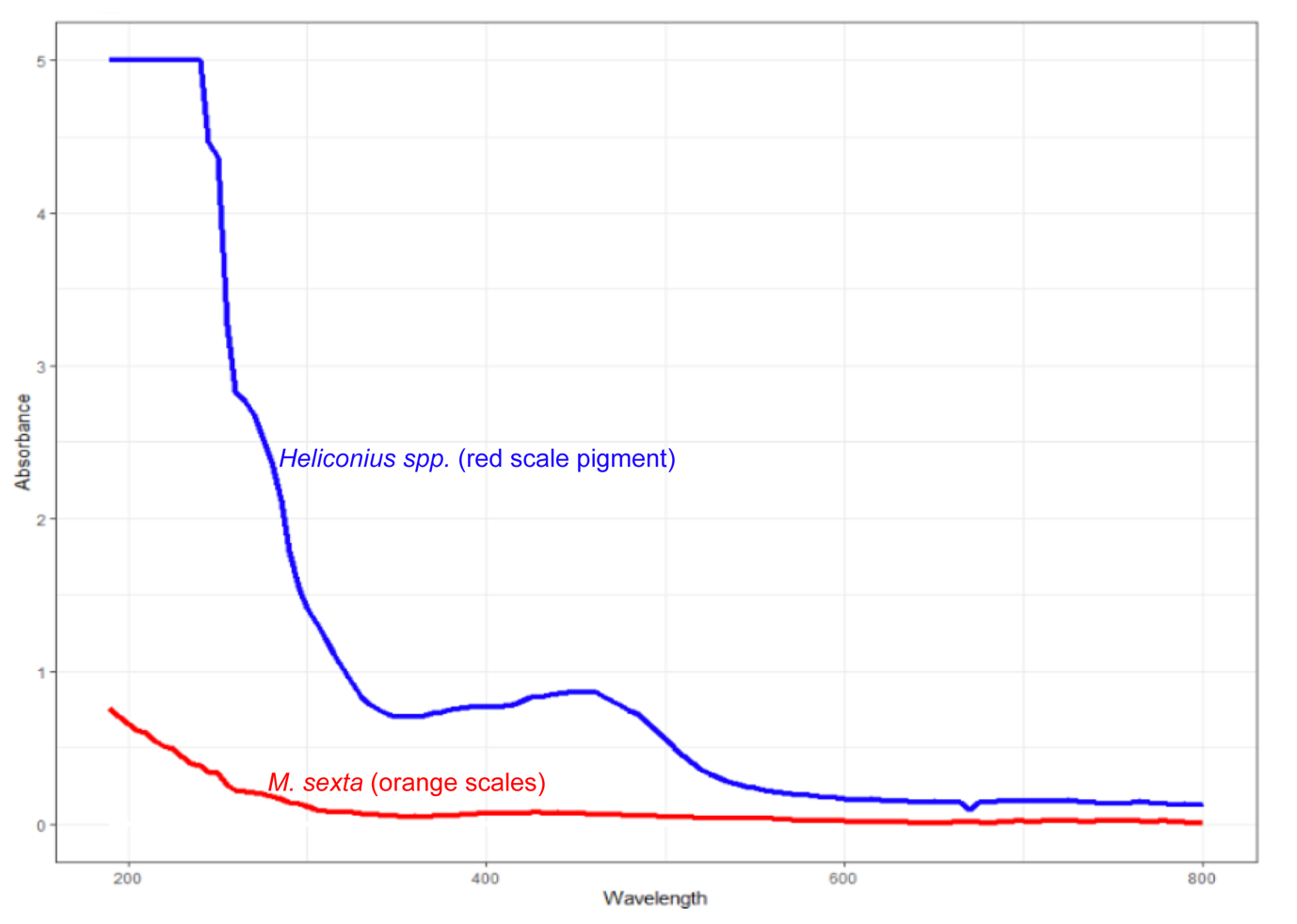

**Figure S2.** ***M. sexta* abdominal pigmentation is insoluble in acid methanol.** UV-Vis absorbance spectra of red ommochrome pigments extracted from *Heliconius* spp. wings (blue line) compared to extracts show a characteristic ommochrome profile (Bybee et al. 2011), while extracts of *M. sexta* orange abdominal scales of (orange line) show almost no detectable absorance in the visual range. This is consistent with pigment insolubility in acidified methanol and suggests a non-ommocrhome chemical identity.

**Table S1. Control micro-injection outcome.**

| **Injection material** | **Number of eggs injected** | **Number of eggs hatched** |
| --- | --- | --- |
| Cas9 (500 ng/µl) | 327 | 229 |
| Nuclease-free water | 207 | 151 |

**Supplementary File 1. Table of RNA-seq data described in Figure 3. Genes highlighted in *yellow* are related to pigmentation pathways. Genes highlighted in *magenta* are known for their role in patterning/segmentation.**
